# Unraveling the gut microbiota of Tibetan chickens: insights into highland adaptation and ecological advantages

**DOI:** 10.1101/2024.02.19.581084

**Authors:** Tao Zeng, Yongqing Cao, Jianmei Yin, Peishi Feng, Yong Tian, Hanxue Sun, Tiantian Gu, Yibo Zong, Xueying Ma, Zelong Zhao, Li Chen, Wenwu Xu, Wei Han, Lizhi Lu

## Abstract

Tibetan animals have several unique advantages owing to the harsh ecological conditions under which they live. Tibetan mammals have been studied extensively. However, understanding of the advantages and underlying mechanisms of the representative high-latitude bird, the Tibetan chicken (*Gallus gallus*, TC), remains limited. The gut microbiota of animals has been conclusively shown to be closely related to both host health and host environmental adaptation. This study aimed to explore the relationships between the cecal microbiome and the advantages of TCs based on comparisons among three populations: native TCs residing on the plateau, domestic TCs living in the plain, and one native plain species. Metatranscriptomic sequencing revealed a significant enrichment of active Bacteroidetes but a loss of active Firmicutes in native TCs. Additionally, the upregulated expression of genes in the cecal microbiome of native TCs showed enriched pathways related to energy metabolism, glycan metabolism, and the immune response. Furthermore, the expression of genes involved in the biosynthesis of short-chain fatty acids (SCFAs) and secondary bile acids (SBAs) was upregulated in the cecal microbiome of native TCs. Data from targeted metabolomics further confirmed elevated levels of certain SCFAs and SBAs in the cecum of native TCs. Based on the multi-omics association analysis, we proposed that the higher ratio of active Bacteroidetes/Firmicutes may be attributed to the efficient energy metabolism and stronger immunological activity of native TCs. Our findings provide a better understanding of the interactions between gut microbiota and highland adaptation, and novel insights into the mechanisms by which Tibetan chickens adapt to the plateau hypoxic environment.

**IMPORTANCE:** The composition and function of the active cecal microbiome were significantly different between the plateau Tibetan chicken population and the plain chicken population. Higher expression genes related to energy metabolism and immune response were found in the cecal microbiome of the plateau Tibetan chicken population. The cecal microbiome in the plateau Tibetan chicken population exhibited higher biosynthesis of short-chain fatty and secondary bile acids, resulting in higher cecal content of these metabolites. The active Bacteroidetes/Firmicutes ratio in the cecal microbiome may contribute to the high-altitude adaptive advantage of the plateau Tibetan chicken population.

## INTRODUCTION

The Tibetan Plateau is the largest uplifted crust on Earth, with an average elevation of 4000 m above sea level. The plateau has a highland continental climate and complex topography with significant variations, rendering it a unique and challenging environment for its inhabitants (1). The harsh ecological conditions in the Tibetan Plateau make it a unique “natural laboratory” for investigating organism adaptations to extreme environments (2). For example, plateau animals have acquired the ability to adapt to extreme cold, low oxygen levels, and strong UV radiation (3). In addition, certain species display distinct behaviors, such as seasonal migration to lower elevations to locate food and seek refuge from inclement weather, as well as burrowing to protect themselves from cold and wind (4). High-altitude organisms have developed effective energy metabolism and immunological capabilities under harsh environmental conditions, enabling them to maintain their health despite harsh external stressors (5, 6). Understanding the mechanisms underlying effective energy conversion and higher stress tolerance in Tibetan animals could potentially contribute to the advancement of the global livestock industry beyond the plateau. However, these mechanisms are not fully understood.

A strong association has been established between gut microbiota and host health as well as host adaptation to the environment (7). Studies conducted on the Tibetan Plateau have revealed the crucial function of gut microbiota in facilitating high-altitude adaptation and health maintenance in both humans and animals in this challenging environment (8). The gut microbiota of native Tibetans and Tibetan pigs varies between high– and low-altitude habitats, indicating the influence of the plateau environment on gut microbiota (5). In addition, a comparative analysis of the gut microbiota in native Tibetan and Han populations residing at different altitudes revealed notable disparities in the compositional and metabolic characteristics of the gut microbial communities. These differences could have implications for health and disease susceptibility (9, 10). In particular, the mutual interaction between host health and gut microbiota is primarily influenced by metabolites produced by symbiotic microorganisms, such as short-chain fatty acids (SCFAs), volatile fatty acids, secondary bile acids (SBAs), and vitamins (11). The role of SCFAs in host immunity has been studied extensively. SCFAs can regulate the epithelial barrier function (12), increase resistance to enteric pathogens (13), and serve as an energy source for colonocyte activity (14). The metabolite succinate, which is primarily derived from dietary fermentation by the gut microbiota, has garnered significant attention. It functions as an intermediate in the TCA cycle, maintains metabolic homeostasis, and acts as a receptor ligand to activate the innate immune response (15). Moreover, SBAs, such as lithocholate (LCA) and deoxycholate (DCA), have been proven to be key innate immune modulators. These metabolites are exclusively produced from primary bile acids (BAs) derived from the host by the symbiotic microbiome (16). Despite numerous valuable findings, research in this field is ongoing, and the intricate relationships between the gut microbiota, health, and highland adaptation should be investigated further.

Tibetan chickens (*Gallus gallus*, TCs) are a unique breed native to the Tibetan Plateau. They have demonstrated remarkable adaptations to high-altitude environments for thousands of years (17, 18). TCs, the most important poultry breed in the Tibetan Plateau, possess several advantages, including excellent disease resistance, medicinal value, and high nutritional content. Moreover, this traditional food is known for its natural and healthy qualities, making it a sought-after product with limited availability and high market price (19). TCs were introduced to plain areas several years ago for large-scale intensive breeding, driven by market demand and economic benefits. However, their introduction into plain areas has resulted in the disappearance of certain valuable traits (17, 20). The gut microbiota of animals is influenced by the microorganisms in their living environment and diet (21). Variations in the cecal microbiome of TCs from five geographic regions have been documented, and the degree of similarity has been correlated with the geographic environment (22). One study found that once TCs were introduced into plain areas, the composition and diversity of the cecal microbiome changed (23). One possible explanation for the physiological changes in TCs living in plains could be alterations in the gut microbiota. Previous studies on the symbiotic microbiomes of TCs have primarily focused on DNA levels using techniques such as 16S rRNA sequencing or metagenomics (24, 25). However, these methods only allow the identification of microorganisms and do not provide information regarding their activity (26). Metatranscriptomics offers valuable insights into microbial gene expression because expressed transcripts serve as surrogates for the actual phenotype (27). With continued advancements in tools and algorithms for analyzing metatranscriptomic data (28), there is an opportunity to delve into the functional activity of the cecal microbiome of TCs.

This study aimed to investigate and compare the active cecal microbiome and its gene expression profiles between chickens residing in plateaus and plains, using metatranscriptomic technology. Additionally, variations in the synthesis capacity of SCFAs and innate immune factors, which are influenced by the cecal microbiome in different chicken populations, were explored. Furthermore, the levels of key metabolites produced by the cecal microbiome, such as SCFAs and BAs, were measured using targeted metabolomics. Through combined analysis of the metatranscriptome and metabolome, our study probed the associations between the unique characteristics of TCs and their cecal microbiome.

## RESULTS

### Active microorganisms and functions in the cecum of chickens

Metatranscriptomics were applied to three chicken populations, yielding an average of 105,456,908 reads per sample. The effective ratio was 99.79%, with 3.17% rRNA (Table S1). The In-TC samples yielded an average of 77,827 contigs, with an N50 length of 899.25 bp and 111,490 ORFs (Table S2). In contrast, the Ex-TC and QY samples had a lower number of contigs, with averages of 11,767 and 7,179, respectively. The N50 lengths for these samples were 735 and 811 bp, respectively (Table S2). The host taxonomy of these transcripts was assigned and compared among the three chicken populations. The results showed that Bacteroidetes and Firmicutes were the dominant active cecal bacterial phyla in chickens residing on the plateau (In-TC) and plain (Ex-TC and QY), respectively (Fig 1A). In addition, the active cecal microbiome had a relatively stable composition in the In-TC samples, whereas significant individual differences were observed in the Ex-TC and QY populations (Fig 1A). These findings were further confirmed by the significantly reduced Bray-Curtis distance of active cecal microbiomes in the In-TC group compared to those in the Ex-TC and QY groups (Tukey’s HSD test, *p* < 0.05, Fig S1). Moreover, comparison of the active cecal bacterial genera among different chicken populations revealed higher activities of *Alistipes*, *Bacteroidetes*, *Desulfovibrio*, *Mediterranea*, and *Prevotella* in the In-TC samples (Tukey’s HSD test, *p* < 0.05, Fig 1B). In contrast, only *Lactobacillus* and *Streptococcus* were more active in the Ex-TC and QY individuals (Tukey’s HSD test, *p* < 0.05, Fig 1B).

**FIG 1.**
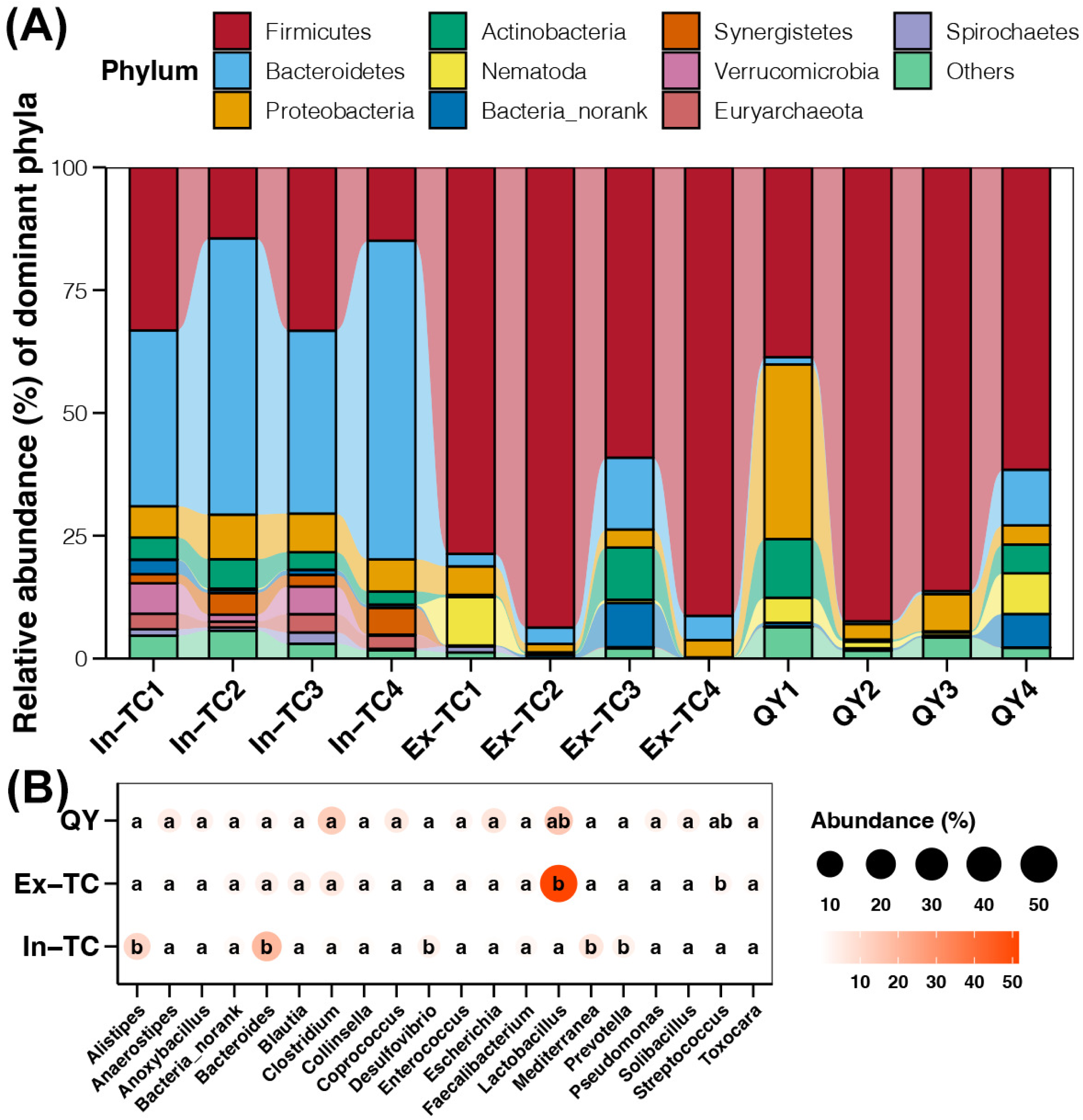
The composition of the active cecal microbiome between the plateau Tibetan chicken population and the plain chicken population. (A) Relative abundance of dominant active cecal bacterial phyla among the samples studied. (B) Differences in the relative abundance of active cecal bacterial genera among the three chicken populations. The size and color of the points represent the average abundance of active cecal bacterial genera in samples from the same population. Different lowercase letters inside the points in the same column represent significant differences (*p*-value of Tukey’s HSD test < 0.05) between different populations.

Based on the comparison of transcripts belonging to different COG terms, the expression levels of genes involved in multiple functions were upregulated in the cecal microbiome of the In-TC group compared to those in the Ex-TC and QY groups (Tukey’s HSD test, *p* < 0.05, Fig 2A). The functions of the In-TC samples included cell division, cell motility, defense mechanisms, energy conversion, signal transduction, and secondary metabolite conversion (Fig 2A). Furthermore, we also observed significantly higher expression of genes related to carbohydrate metabolism in the cecal microbiomes of In-TC samples compared to those of the Ex-TC and QY groups (Tukey’s HSD test, *p* < 0.05, Fig 2B).

**FIG 2.**
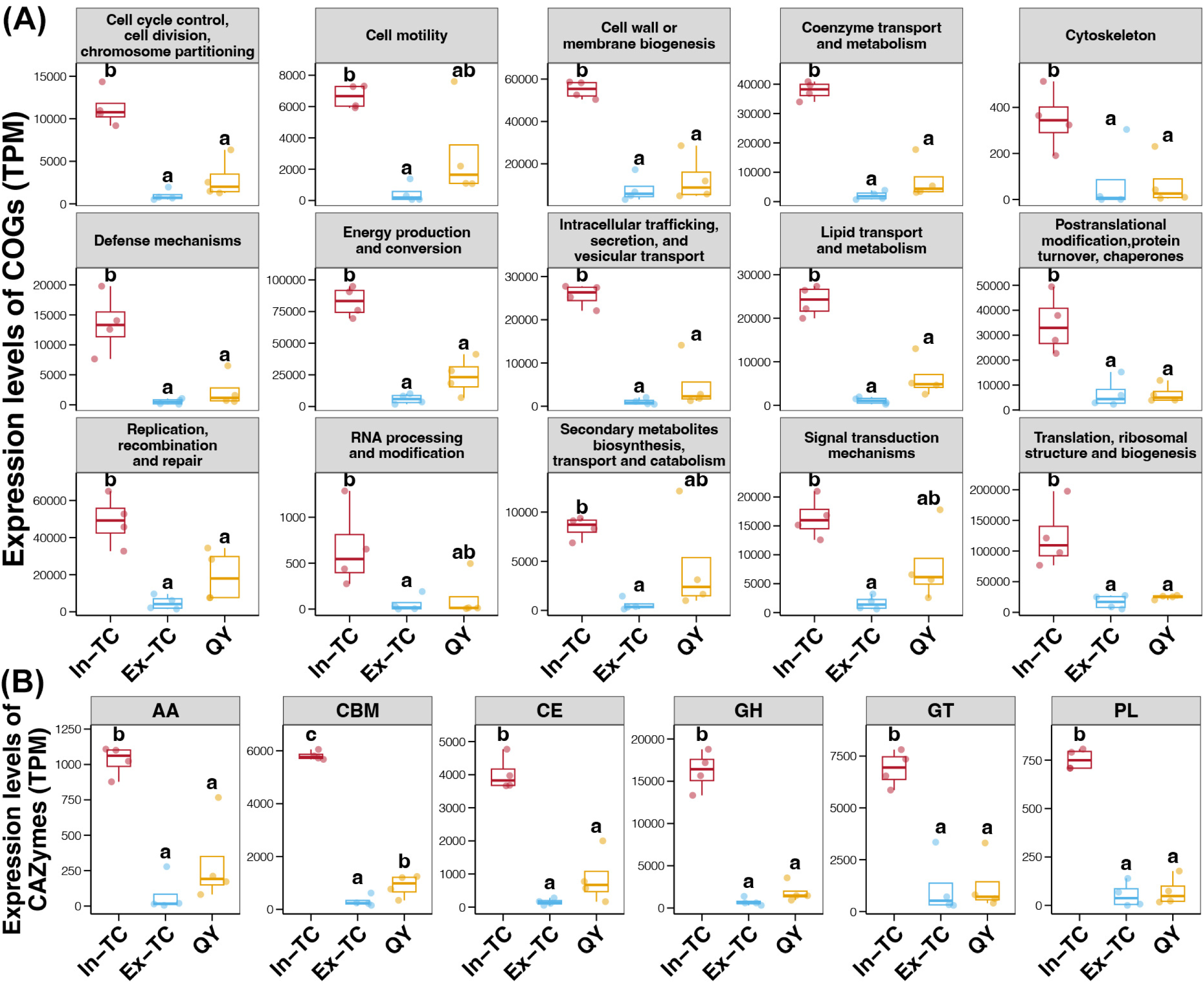
Expression levels of COG terms and CAZymes in the cecal microbiome between the plateau Tibetan chicken population and the plain chicken population. Differences in the expression levels of COG terms (A) and CAZymes (B) in the cecal microbiome of the three chicken populations. Different lowercase letters above the boxes in the same subFig represent significant differences (*p*-value of Tukey’s HSD test < 0.05) between different populations.

### Identification of DEGs and their functions in the cecal microbiome of chicken populations

The PCoA plot revealed distinct clusters of gene expression profiles between the cecal microbiomes of chickens residing in the plateau (In-TC) and plain (Ex-TC and QY) regions (Fig S2A). The Adonis test also confirmed significant differences in the gene expression profiles of the cecal microbiomes among the three chicken populations studied (*p* < 0.05). The In-TC populations had a total of 95,455 and 96,014 DEGs in their cecal microbiomes compared to the Ex-TC and QY populations, respectively (Fig S2B and S2C). Most of these DEGs (92,347 and 90,267, respectively; Fig S2B and S2C) were upregulated in the In-TC group, accounting for over 95% of the DEGs. In contrast, only 6151 genes were found to be differentially expressed in the cecal microbiomes between the Ex-TC and QY populations (Fig S2D). The number of DEGs in the comparison of the Ex-TC and QY groups significantly decreased, primarily because of the decrease in upregulated genes (2635 compared to more than 90,000).

Considering those chickens from the Ex-TC and QY populations both lived in plain environments, we identified potentially critical genes for the high-latitude adaptability of TCs by examining DEGs that were present in the comparisons of In-TC with Ex-TC or QY, but not in the comparison of Ex-TC with QY. Based on Venn diagrams, we identified 89,971 upregulated and 440 downregulated genes that were differentially expressed in the cecal microbiomes of chickens residing in the plateau and plain regions (Fig 3A and 3B). The upregulated genes were enriched in several KEGG pathways related to energy metabolism, glycan metabolism, and immune response (Fig 3C). No enriched pathways were identified for the downregulated genes.

**FIG 3.**
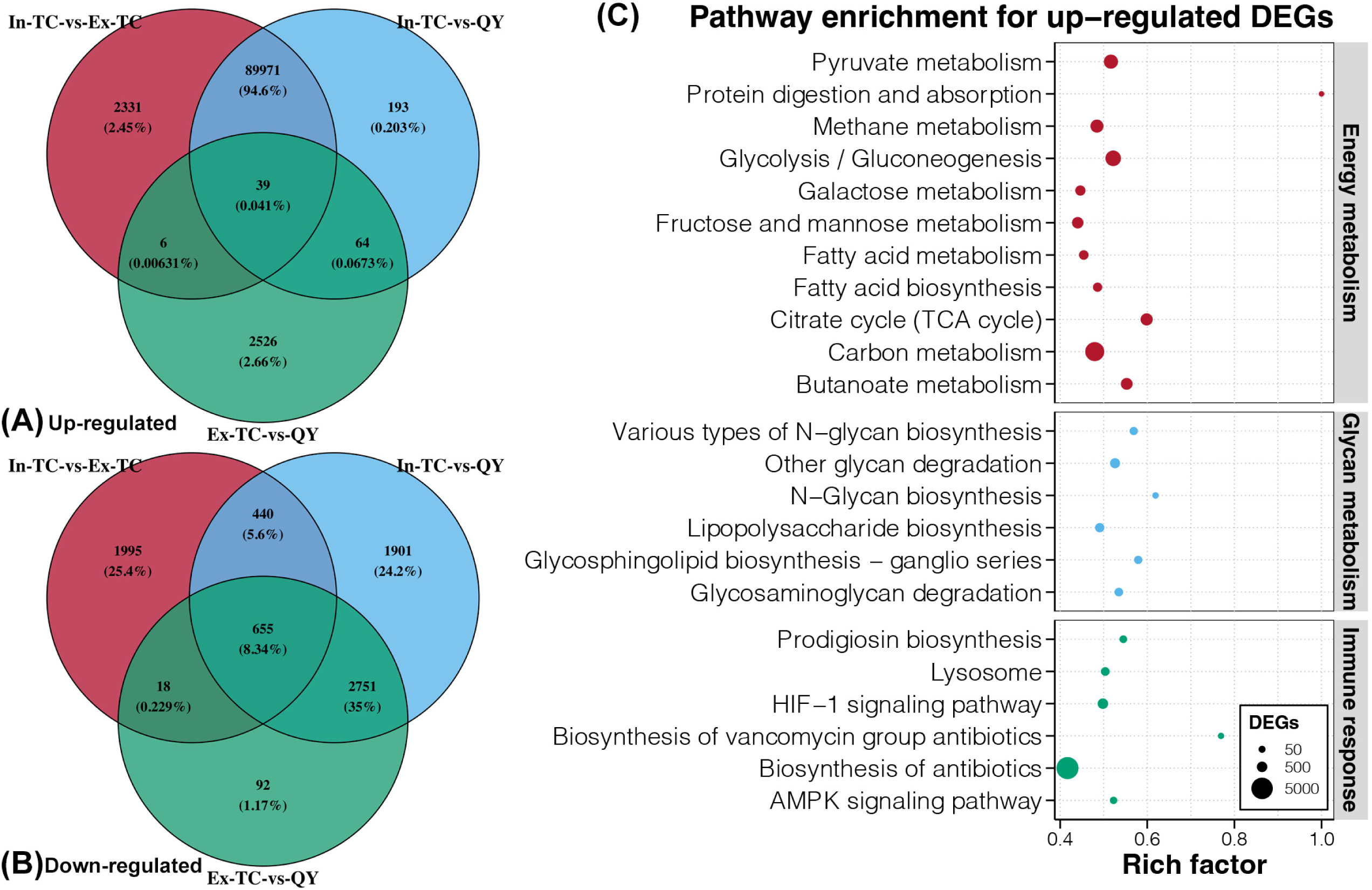
DGEs and enrichment analysis in the cecal microbiomes between the plateau Tibetan chicken population and the plain chicken population. Venn diagrams for upregulated (A) and downregulated (B) DEGs among the three comparisons. (B) KEGG enrichment analysis of upregulated genes in the cecal microbiomes of chickens residing in the plateau and plain regions.

### Powerful biosynthesis of SCFAs and immune activity in the cecal microbiome of In-TCs

Regarding genes responsive to the synthesis of SCFAs, *pct* and *ACSS1_2* for acetate and propionate biosynthesis, respectively, and *buk* and *atoAD* for butyrate biosynthesis showed significantly higher expression in the cecal microbiome of In-TC individuals than in Ex-TC and QY samples (Fig 4A). The hosts of the *pct* gene in all three populations belonged to the phylum Firmicutes (Fig 4B), indicating that discrepancies in gene expression could be attributable to host differences at the fine taxonomic level. Further research revealed that the *Anaerotignum* genus was the primary host of the *pct* gene in In-TC samples, whereas the *Ruminococcus* genus was the primary host in Ex-TC and QY samples (Fig S3). In all three populations, the host of the *ACSS1_2* gene was the *Desulfovibrio* genus belonging to the phylum *Proteobacteria* (Fig 4B and Fig S4). The abundance of active *Desulfovibrio* was significantly higher in the In-TC samples than in the Ex-TC and QY samples (Fig 1B). This finding was consistent with the upregulated expression of the *ACSS1_2* gene in In-TC individuals. The hosts of *buk* and *atoAD* genes in the studied samples were mostly from the Bacteroidetes phylum (Fig 4B), particularly in the In-TC samples (100%). The higher expression levels of *buk* and *atoAD* genes in the In-TC population were consistent with the higher abundance of active Bacteroidetes in the cecal microbiome of In-TC individuals (Fig 1A).

**FIG 4.**
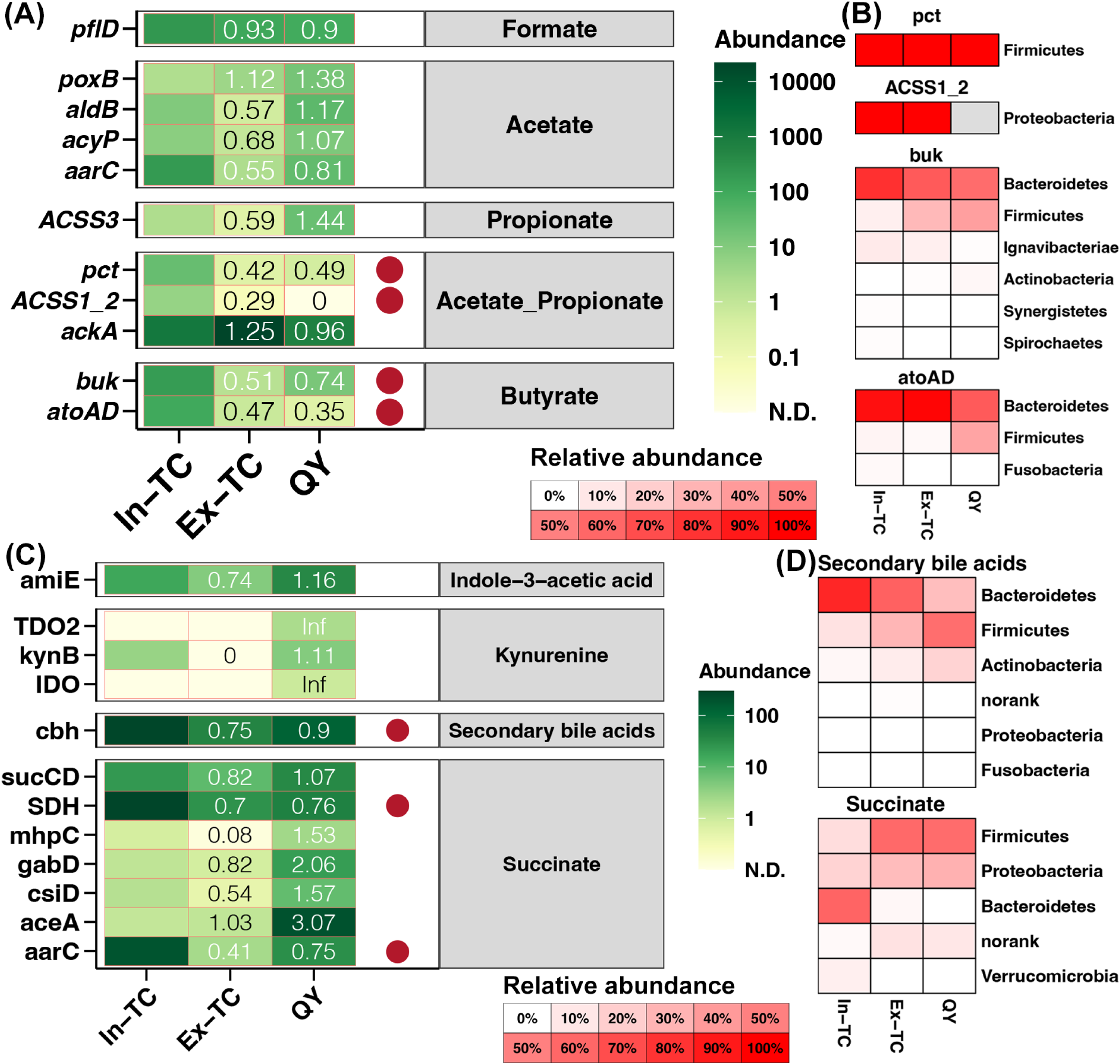
Biosynthesis of SCFAs and immune activity in the cecal microbiome of In-TCs. (A) Expression levels of genes associated with SCFA biosynthesis in the cecal microbiome of different chicken populations. (B) Host composition of SCFA biosynthesis genes differentially expressed among different chicken populations. (C) Expression levels of genes related to the biosynthesis of innate immune factors in the cecal microbiome of different chicken populations. (D) Host composition of innate immune factor biosynthesis-related genes differentially expressed among different chicken populations. Color and values represent the fold change of gene expression levels in each population compared to that in the In-TC group. Red points represent significant differences in gene expression levels among the different groups.

The expression levels of several genes involved in the biosynthesis of innate immune factors were further extracted from metatranscriptome data and compared across different populations. Secondary bile acid biosynthesis gene, *cbh*, and succinate synthesis genes, *SDH* and *aarC,* were found to be significantly upregulated in the cecal microbiome of In-TC individuals compared to Ex-TC and QY samples (Fig 4C). In the In-TC populations, the Bacteroidetes phylum was the primary host for these three genes, whereas the Firmicutes phylum was the primary host in the Ex-TC and QY groups (Fig 4D). These findings corroborated the increased activity of Bacteroidetes and decreased activity of Firmicutes in the cecal microbiome of In-TC individuals (Fig 1A).

### Correlations between the cecal microbiome and metabolome in chickens

Target metabolomics data revealed that the cecum of different chicken populations displayed distinct compositions of SCFAs and Bas, as indicated by the PCoA and adonis test (Fig 5A and 5B). In addition, the cecum of In-TC individuals exhibited a significantly higher concentration of acetic acid than that of the Ex-TC and QY groups (Fig 5C). Moreover, the cecum of In-TC individuals showed a higher abundance of multiple BAs than the Ex-TC and QY populations, including UDCA, UCA, isoLCA, GUDCA, GLCA, GHDCA, GDCA, beta-UDCA, beta-MCA, and 7-ketoLCA (Fig 5C). In contrast, only norDCA decreased in the cecum of In-TC individuals (Fig 5C).

**FIG 5.**
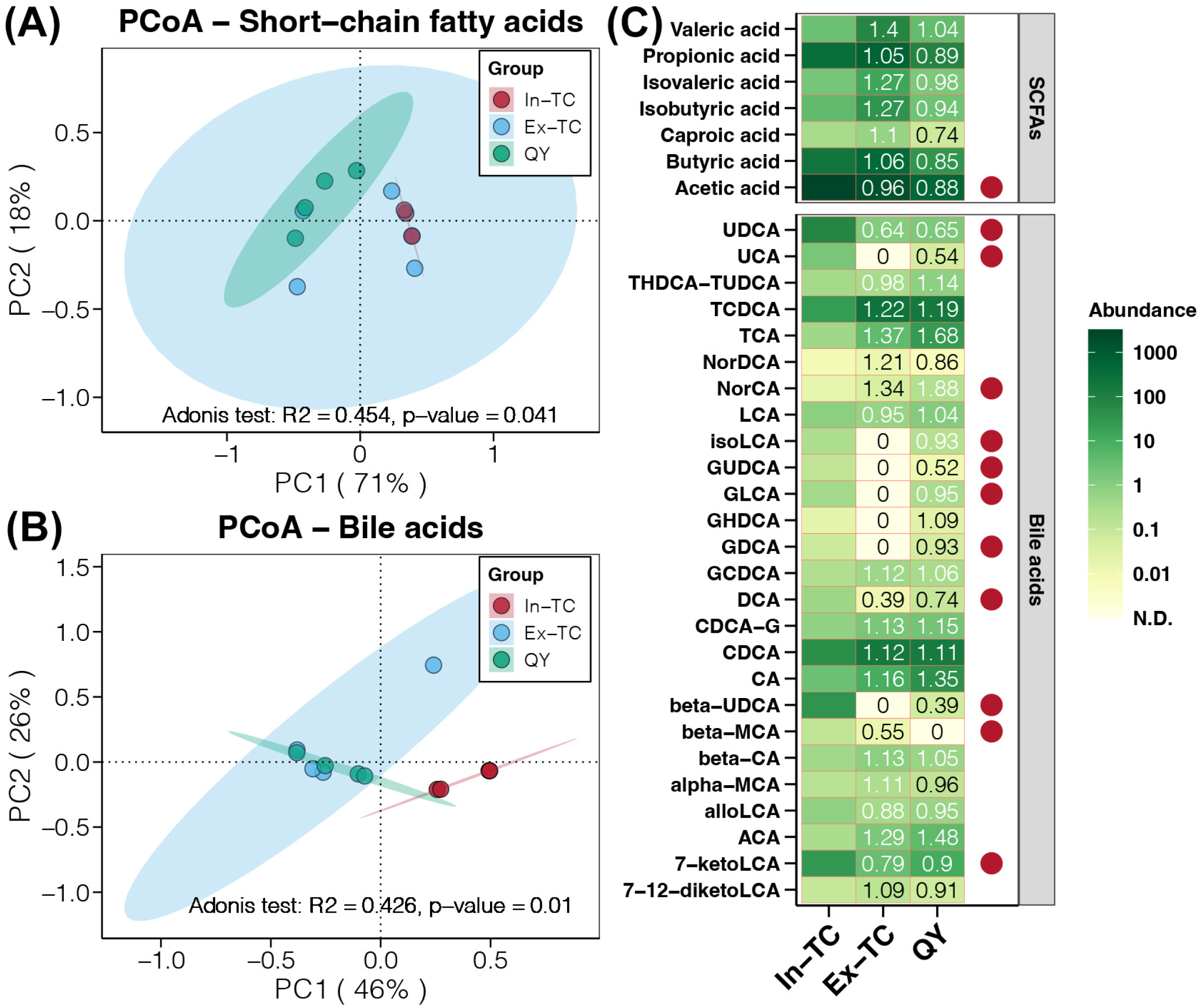
Targeted metabolomic analysis of SCFAs and BAs in the cecum among the three chicken populations. PCoA showing differences in the composition of SCFAs (a) and BAs (b) in the cecum among the three chicken populations studied. (c) Comparison of SCFA and BA concentrations in the cecum of different chicken populations. Colors and values represent the fold-change in metabolites in each population compared with those in the In-TC group. The red points represent significant differences in metabolite concentrations among the different groups.

The above results indicated that the abundance of Bacteroidetes and Firmicutes in the cecal microbiome could contribute to the high-altitude adaptive advantage of TCs. The ratio of Bacteroidetes/Firmicutes in the gut microbiota has been proven to be an indicator of health in humans and rats (29, 30). In this study, we analyzed the ratio of active Bacteroidetes/Firmicutes in the cecal microbiome of chickens and compared them between populations residing in the plateau and plain regions. A significantly higher ratio of active Bacteroidetes/Firmicutes was observed in the cecal microbiome of In-TC individuals than that in the Ex-TC and QY samples (Tukey’s HSD test, *p* < 0.05, Fig 6). In all analyzed samples, we found significant positive correlations between the ratio of active Bacteroidetes/Firmicutes and the expression levels of SCFA biosynthesis genes (*pct*, *atoAD*, and *buk*) (Fig 6). They were further positively correlated with propionic acid, acetic acid, and butyric acid contents in the cecum of chickens (Fig 6). Furthermore, significant positive correlations were observed between the ratio of active Bacteroidetes/Firmicutes and expression levels of genes involved in the biosynthesis of innate immune factors (*SDH*, *aarC*, and *cbh*) (Fig 6). These genes were positively correlated with the content of diverse BAs in the cecum of chickens (Fig 6).

**FIG 6.**
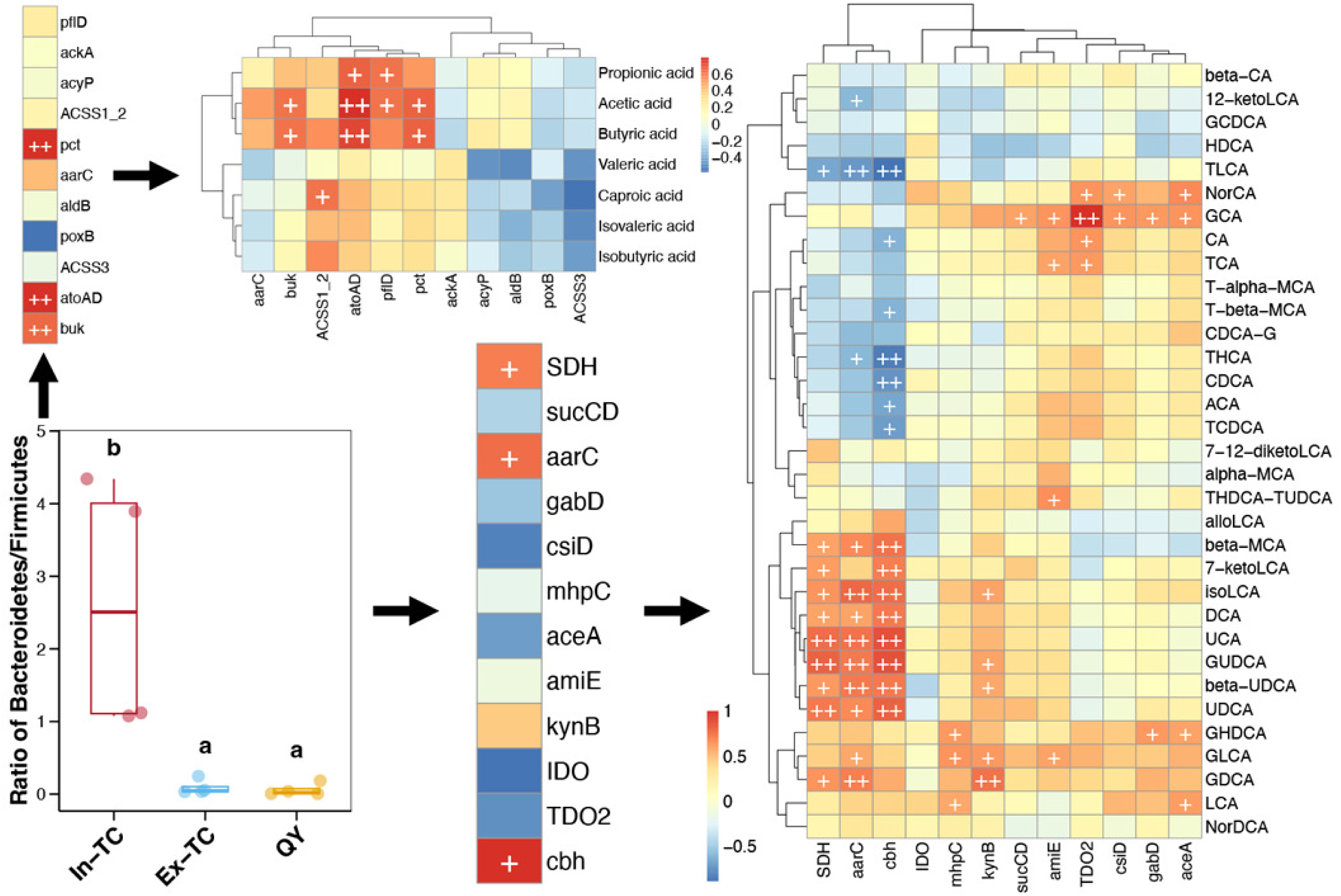
Differences in the ratio of active Bacteroidetes/Firmicutes in the cecal microbiome between chickens residing in plateau and plain areas and their correlations with gene expression levels and metabolite contents related to SCFAs and BAs. Different lowercase letters above the boxes in the boxplot represent statistically significant differences (*p*-value of Tukey’s HSD test < 0.05) between different populations. The color of the blocks in the heatmaps represents the correlation coefficient of Spearman correlations. “+” and “++” represent p-values Spearman correlations lower than 0.05 and 0.01, respectively.

## DISCUSSION

This is the first study to use metatranscriptomic technology to investigate the active microorganisms and their physiological functions in the cecum of TCs. We observed significant variations in the composition and function of the active cecal microbiome between chickens residing in the plateau and plain regions, based on comparisons between the In-TC, Ex-TC, and QY populations. In contrast, the active cecal microbiome of TCs became remarkably similar to that of other plain populations once introduced into the plains. Previous studies have indicated that both genetic and environmental factors influence the symbiotic microbiome of chickens (31, 32). In the present study, the In-TC and Ex-TC populations had similar genetic backgrounds. Thus, the differences in the active cecal microbiome between the two groups could be attributed to habitat factors, including environmental conditions and dietary differences. Given that the Ex-TC and QY populations in our study resided on the same farm, shared a consistent diet, and were exposed to similar environmental factors, the presence of a similar active cecal microbiome in both groups highlights the significant influence of habitat on genetic constraints. These findings indicate that maintaining the advantages related to the cecal microbiome in TCs became challenging once they were introduced into plain areas.

At the taxonomic level, *Alistipes*, an active cecal bacterial genus abundant in the In-TC population, has been associated with potential benefits in protecting the human gastrointestinal tract against colitis, autism, and cirrhosis (33). In addition, another In-TC-enriched active cecal bacterial genus, *Prevotella*, has been found to be more abundant in Tibetan animals, which benefits the host by maintaining gut energy homeostasis and adapting to hypoxic pressure (34, 35). *Streptococcus*, an active cecal bacterial genus enriched in chicken populations residing in plains, is a pathogen that causes various infections in humans (36). Streptococcal bacteria are highly contagious and can easily spread through droplets produced when an infected person coughs or sneezes, as well as through shared food or drinks (37). At the functional level, the expression levels of all carbohydrate-active enzymes were higher in the In-TC population. These enzymes degrade carbohydrates in food to generate energy (38). The upregulated DEGs in the In-TC cecal microbiome were enriched in various energy metabolism-related pathways. These pathways include fatty acid and saccharide metabolism as well as the TCA cycle. These findings indicate that TCs residing on the plateau use energy more efficiently and pose relatively low human risks.

*Alistipes* species, in addition to their role in disease protection, are indole-positive, which affects the serotonin precursor tryptophan (33). These factors are closely linked to host innate immunity (11). Several glycan-related pathways were enriched in the active cecal microbiome of the In-TC population. Glycans are complex carbohydrates that play crucial roles in the regulation of various aspects of human physiology and pathology. These functions include signal transduction and defense against microbial infections (39). Glycans, in particular, act as molecular patterns recognized by glycan-binding proteins and hence play a crucial role in various cellular processes, including cell-cell interactions and immunological responses (40). Additionally, we observed that some immune-related pathways were enriched in the active cecal microbiome of the In-TC population, including pathways for immunological signal transduction (AMPK and HIF-1) and biosynthesis of antibiotics that can effectively eliminate pathogens (prodigiosin and vancomycin). Furthermore, our findings revealed higher contents of acetic acid and several SBAs in the cecum of In-TC individuals. Acetate, an SCFA, is produced by symbiotic microorganisms and plays a crucial role in maintaining the overall health of the host (41). Acetate may affect gut health by improving intestinal barrier function via cross-feeding mechanisms (42). Acetate has also been associated with epithelial maintenance, wound healing, and improved barrier function in the gastrointestinal tract. Research has demonstrated that ursodeoxycholic acid (UDCA) regulates the defense mechanisms of antimicrobial peptides, protects epithelial cells, and exerts anti-inflammatory effects in the cecum of the In-TC population (43). Another enriched SBA, isoLCA (isolithocholic acid), has been found to be one of the most abundant BAs in the gastrointestinal tract and is regarded as an essential modulator of immunological homeostasis (44). These results suggest that a higher level of immunological activity is mediated by the cecal microbiome in TCs residing on the plateau.

Our study combined metatranscriptomics and targeted metabolomics technologies to explore the underlying mechanisms of the active cecal microbiome that contribute to the advantages of TCs living in the Tibetan Plateau. Association analysis yielded a framework that could explain the higher immunological activity of TCs residing on the plateau. The ratio of active Bacteroidetes/Firmicutes increases in the cecal microbiome of TCs residing in the plateau, which induces the powerful biosynthesis of SCFAs and SBAs to support increased immunological activity. In contrast, the ratio of active Bacteroidetes/Firmicutes was lower in TCs and QYs living in the plain, resulting in reduced SCFA and BA content in the cecum. Notably, a recent analysis of the gut microbiota of Tibetan Plateau animals revealed contradictory results to our findings, with the authors obtaining a lower Bacteroidetes/Firmicutes ratio in higher-altitude mammals than in lower-altitude mammals (8). As the dominant phyla of the mammalian gut microbiota, the relative abundance of Bacteroidetes and Firmicutes can serve as indicators of obesity, with a higher abundance of Firmicutes and a lower abundance of Bacteroidetes in obese individuals (29, 45). This is consistent with the fact that high-altitude animals require additional body fat to resist cold. However, the higher ratio of active Bacteroidetes/Firmicutes observed in the In-TC samples contradicts previous findings. One possible explanation for this conflict could be the variation in the research methods. Our study measured active microorganisms based on microbial RNA sequencing, which differs from previous studies that relied on microbial DNA. Another potential explanation could be the physiological differences between mammals and birds. Previous research in Arbor Acres broiler chickens indicated that, unlike in mammals, the Bacteroidetes/Firmicutes ratio in birds cannot be used as an indicator of gut health (46). Our findings suggest that the presence of active Bacteroidetes/Firmicutes could serve as a significant marker of innate immunological activity mediated by the cecal microbiome in chickens. However, further large-scale population comparisons and validation experiments are necessary to confirm the accuracy and feasibility of regulating active Bacteroidetes/Firmicutes to improve immunity in chickens.

## CONCLUSION

In this study, we present an initial investigation of the active cecal microbiome and its potential advantages in plateau-dwelling TCs over plain-dwelling TCs. The composition and function of the active cecal microbiome vary significantly across different chicken populations. In particular, the higher expression of functional genes in the cecal microbiome suggests that TCs residing on the plateau have more effective energy metabolism and a stronger innate immune response than individuals residing on the plains. Moreover, the cecal microbiome in the native TC population has an increased ability to produce SCFAs and SBAs, resulting in higher cecal content of these metabolites. Finally, the advantages of native TCs are linked to the active Bacteroidetes/Firmicutes ratio in the cecal microbiome. Our findings deepen the understanding of the interactions between gut microbiota and highland adaptation and provide valuable insights into the mechanisms by which Tibetan chickens adapt to the hypoxic environment of the plateau.

## METHODS

### Study animals and sample collection

Two different chicken species, TCs and Qingyuan (QY) chickens, were used in our study. Two different populations were selected from the TC individuals. One population included TCs that always resided on the Tibetan Plateau after birth (In-TC), and the other was a transitional TC population residing in plain environments ex situ over 20 years (Ex-TC). The QY species was a native breed in plain environments. The Ex-TC and QY populations were raised in the same environment to exclude the influence of diet and breeding environment. This sample setting considered both genetic and environmental variations in the population, enabling a more accurate assessment of the uniqueness of TCs. Twelve individuals were selected from each population for sample collection. All study animals were approximately 300 days old, the chickens were healthy, and had not received any medications for at least three months prior to sampling. The cecal content of each individual was collected under sterile conditions. Subsequently, the contents of three individuals from the same population were combined and added to 1.5 ml sterile polypropylene tubes. All samples were immediately snap-frozen in liquid nitrogen until further analysis.

### RNA extraction and metatranscriptomic sequencing

Total RNA was extracted from the cecal contents using TRIzol Reagent (Invitrogen, Carlsbad, USA), and genomic DNA was removed using DNase I (TAKARA, Dalian, China). Next, the quality of the extracted RNA was assessed using an Agilent 2100 Bioanalyzer and NanoDrop ND-2000. Metatranscriptome libraries (450 bp) were constructed using the TruSeqTM Stranded Total RNA Sample Preparation Kit (Illumina, San Diego, CA, USA) and sequenced on the Illumina NovaSeq 6000 platform with a pair-end 150 bp strategy at Shanghai BIOZERON Co., Ltd. (Shanghai, China). The raw reads underwent quality control using Trimmomatic with a sliding window (4:15) and minlen (75) settings. Next, the clean reads were aligned to the chicken reference genome, and the mapped reads were removed. The rRNA reads from the clean reads were further removed by alignment to the SILVA database using SortMeRNA software (47). The MegaHit algorithm (48) was used to assemble the remaining mRNA reads into contigs, and open reading frames were predicted using METAProdigal (49). The CD-HIT method (50) was used to remove redundancy and construct a non-redundant gene catalog with 95% identity and 90% coverage. All unique genes were searched against the eggNOG (51), CAZyme (52), and KEGG (53) databases for functional annotation using BLASTp with a cut-off E-value less than 1e-5. The genes were further aligned to the NR database using BLASTp to assign host taxonomy. The expression level of each gene in each sample was calculated using Salmon (54). Three comparisons were conducted to identify differentially expressed genes (DEGs): In-TC vs. Ex-TC, In-TC vs. QY, and Ex-TC vs. QY. Based on the results obtained from the “edgeR” package (55), genes with a log-trans fold change greater than 2 and a p-value adjusted by the false discovery rate method less than 0.5 in each comparison, were recognized as DEGs.

The biosynthesis-related functional analyses in this study primarily focused on SCFAs and certain innate immune factors generated by microbes. Specifically, SCFAs include formate, acetate, propionate, and butyrate. Additionally, innate immune factors include indole-3-acetic acid (IAA), indole-3-aldehyde (IAId), kynurenine, SBAs, succinate, and trimethylamine N-oxide (TMAO). The genetic potential of the gut microbiota was analyzed by sorting marker genes based on the KEGG database. The detailed marker genes and abundance calculation processes used in this study are listed in Supplementary Data 1. Based on the above KEGG annotations, nucleotide sequences of genes encoding the selected marker genes were searched against the NCBI NR database using BLASTX. The top 50 hits with E-values < 0.001 were imported into MEGAN, and the taxonomy was assigned using the lowest common ancestor algorithm.

### Targeted metabolomic analysis

Standards for SCFAs and BAs were obtained from Sigma-Aldrich (St. Louis, MO, USA). Targeted metabolome analysis involved resuspending the cecum digesta in liquid nitrogen and diluting the sample by vigorous vortexing in water. Next, a 50 μL fraction was homogenized with 200 μL of acetonitrile/methanol (1:1) containing the appropriate internal standards. The mixture was vortexed thoroughly to ensure homogenization. The mixture was then placed on ice for 30 min. The mixture was then centrifuged at 12,000 rpm for 10 min. An ultra-high-performance liquid chromatography coupled with tandem mass spectrometry (UHPLC-MS/MS) system (ExionLC™ AD UHPLC-QTRAP 6500+, AB SCIEX Corp., Boston, MA, USA) was used to quantify the SCFAs and AAs. Separation was carried out on an ACQUITY UPLC BEH Amide column (2.1 × 100 mm, 1.7 μm) that was maintained at 50 °C. The mobile phase, consisting of 0.1% formic acid in 5 mM ammonium acetate (solvent A) and 0.1% formic acid in acetonitrile (solvent B), was delivered at a flow rate of 0.30 mL/min. The solvent gradient was set as follows: initial 80% B, 0.5 min; 80-70% B, 2 min; 70-45% B, 4 min; 45-80% B, 6.01 min; 80% B, 9 min. The mass spectrometer was operated in the positive multiple reaction mode (MRM). The parameters used were as follows: IonSpray Voltage (5500 V), Curtain Gas (35 psi), Ion Source Temperature (550 °C), and Ion Source Gas of 1 and 2 (50 and 60 psi, respectively).

### Statistical analysis

All statistical analyses were conducted using the R v4.2.2 platform and the results were visualized using the “ggplot2” package. Variations in the abundance of active cecal microorganisms, COG terms, and CAZyme categories in cecal microbiomes among the three chicken populations were analyzed using Tukey’s HSD test (“multcomp” package). PCoA and adonis test based on the Bray-Curtis distance (“vegan” package) was performed to explore the differences in the composition of and gene expression profiles of active cecal microbiome as well as the compositions of SCFAs and BAs in the cecum across three different chicken populations. A Venn diagram (“VennDiagram” package) was used to identify the potential key genes related to the high-altitude adaptability of TCs based on the recognized DEGs. KEGG enrichment analysis of these putative key genes was performed using the KOBAS tool (56). Differences in the expression levels of genes related to SCFA biosynthesis and innate immune factors in the cecal microbiomes and the contents of SCFAs and BAs in the cecum among the three chicken populations were also compared using Tukey’s HSD test. Tukey’s HSD test was also used to evaluate variations in the ratio of active Bacteroidetes/Firmicutes in the cecal microbiomes among the three chicken populations. The correlations between the ratio of active Bacteroidetes/Firmicutes, gene expression levels related to SCFA biosynthesis and innate immune factors, and the contents of SCFAs and BAs were assessed using Spearman correlation (“psych” package).

## SUPPORTING INFORMATION

**Table S1.** Statistics of metatranscriptome sequencing.

**Table S2.** Statistics of assembly and gene recognition.

**Fig S1.** Differences in the inter-individual variations of active microbiome among different populations based on the Bray-Curtis distance.

**Fig S2.** Identification of DEGs in the cecal microbiome of chicken populations. (A) PCoA illustrating the differences in gene expression profiles of the cecal microbiome among the three studied chicken populations. Volcano plots were used to identify the DEGs in cecal microbiome in the comparisons of In-TC vs. Ex-TC (B), In-TC vs. QY (C), and Ex-TC vs. QY (D), respectively.

**Fig S3.** Host composition of the pct gene in cecal microbiome of chickens.

**Fig S4.** Host composition of the *ACSS1_2* gene in cecal microbiome of chickens. **Supplementary Data 1.** Key genes for biosynthesis of nutrients and innate immune factors in gut microbiota of animals.

## AUTHOR CONTRIBUTIONS

Tao Zeng, Lizhi Lu and Wei Han planned the project. Tao Zeng, Lizhi Lu, Wei Han, Yong Tian and Peishi Feng designed the research. Tao Zeng, Yongqing Cao, Jianmei Yin, Hanxue Sun, Tiantian Gu, Xueying Ma and Li Chen prepared the research materials. Tao Zeng, Yongqing Cao, Yibo Zong, Zelong Zhao and Wenwu Xu performed the experiments for sequencing and the data analyses. Tao Zeng drafted the manuscript. Lizhi Lu and Wei Han revised the manuscript. All authors read and approved the final manuscript.

## ACKNOWLEDGMENTS

This work was supported by National Key Research and Development Program of China (2021YFD1200302), Key Research and Development Program of Zhejiang Province (2021C02034), Project of Key Research and Development Plan (Modern Agriculture) of Jiangsu Province (BE2019353) and “JBGS” Project of Seed Industry Revitalization in Jiangsu Province (JBGS [2021] 029).

## DATA AVAILABILITY STATEMENT

The raw reads of metatranscriptomic sequencing for all samples have been deposited in the NCBI Sequence Read Archive (SRA) under the accession number of PRJNA1060358. Targeted metabolomics data of SCFAs and Bas have been deposited in the OMIX, China National Center for Bioinformation / Beijing Institute of Genomics, Chinese Academy of Sciences (https://ngdc.cncb.ac.cn/omix: accession no.OMIX 005604 and OMIX 005605).

## ETHICS STATEMENT

Animals used in this study were raised in accordance with the national standard of Laboratory Animal Guidelines for ethical review of animal welfare. All experiment procedures were approved by the Animal Use Committee of Zhejiang Academy of Agricultural Sciences (No. 20-022).

---

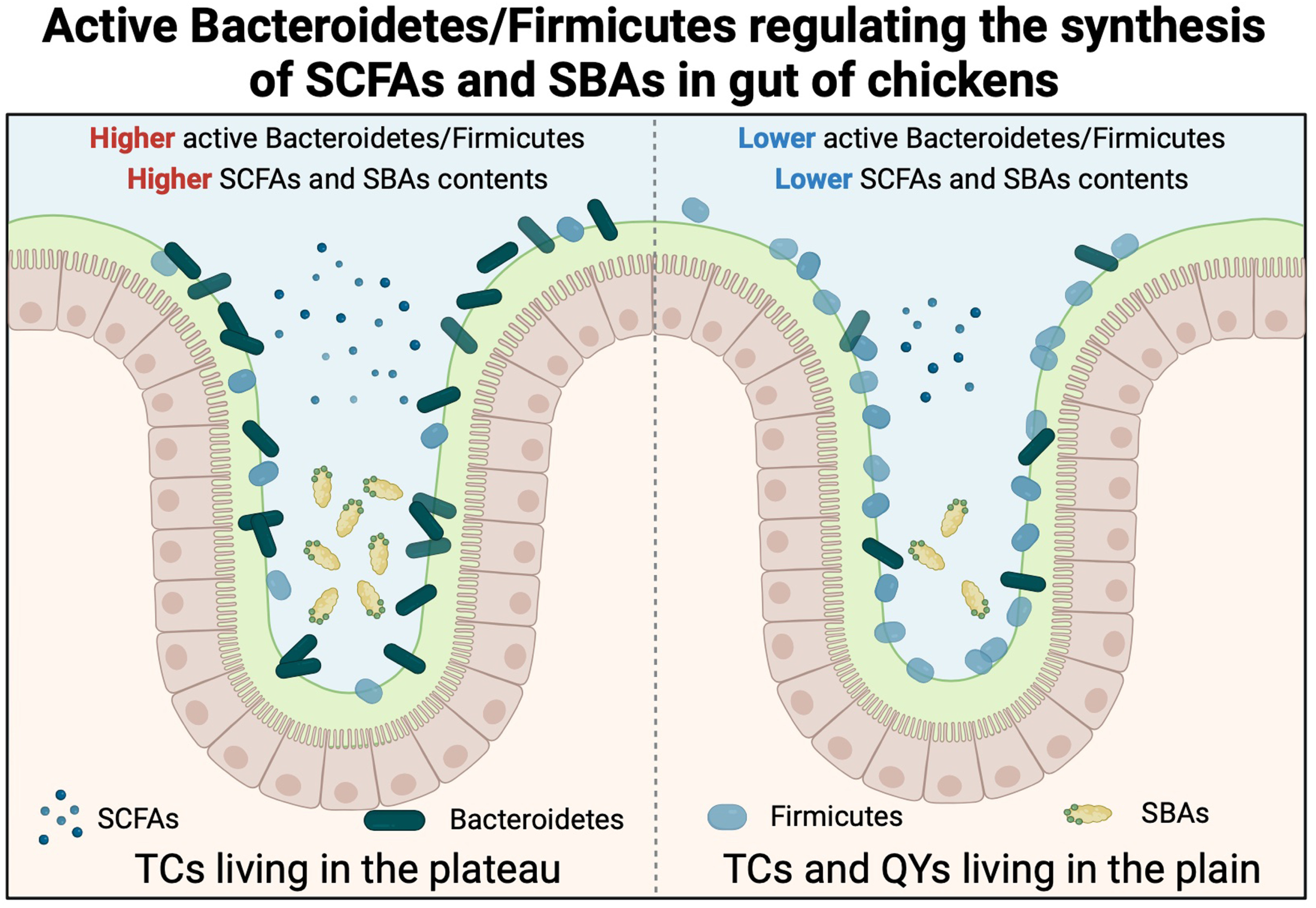

